# The unique properties of Big tau in the visual system

**DOI:** 10.1101/2023.07.11.548562

**Authors:** Itzhak Fischer, Theresa Connors, Julien Bouyer, Ying Jin

## Abstract

Tau is a microtubule associated protein that plays important roles in regulating the properties of microtubules and axonal transport, as well as tauopathies associated with toxic aggregates leading to neurodegenerative diseases. It is encoded by the MAPT gene forming multiple isoforms by alternative splicing of exons 2/3 at the N-terminal and exon 10 which determines the numbers of microtubule binding repeats (3R or 4R). The high molecular weight (MW) tau isoform termed Big tau contains an additional large exon 4a generating a long projecting domain and expressed as a 110 kDa protein. Big tau was originally discovered in the peripheral nervous system but later found in selective CNS areas that project to the periphery as well as in the cerebellum and the visual system. However, there is a gap of knowledge in understanding the expression patterns and the role of Big tau during normal neuronal development and pathological conditions relative to the common low MW isoforms. Here we investigated the properties of Big tau in the retina and optic nerve and in particular its unique isoform structure as a middle MW of 90kDa and its distribution in retinal ganglion cells and axons of the optic nerve. We discovered that Big tau expresses the 4a exon as well as exons 6 and 10 (4R), lacking exons 2/3 but sharing the extensive phosphorylation characteristic of other tau isoforms. Importantly, early in development the visual system expresses only the low MW isoform (3R) switching to both the low and middle MW isoforms (4R) in adult retinal ganglion neurons and their corresponding axons. This is a unique structure and expression pattern of Big tau likely associated with different properties than what has been previously described, requiring more research to elucidate the detailed roles of Big tau in the visual system.

## 1. Introduction

### 1.1. What is Big tau

Tau is a microtubule associated protein (MAP) that modulates the dynamic properties and organization of microtubules in neurons as well as axonal transport, interactions with other structural proteins and various signaling pathways (Wang & Mandelkow, 2016). Tau can also form insoluble toxic aggregates and fibrils associated with neurodegenerative diseases (e.g., Alzheimer’s disease) and tauopathies (e.g., frontotemporal dementia) (Lee & Leugers, 2012). Tau is encoded by the MAPT gene and shows remarkable heterogeneity, and as the result of alternative splicing generates six developmentally regulated isoforms of 45-60 kDa. These isoforms represent a combination of different numbers of microtubule binding repeats (3R or 4R) by selective expression of exon 10, and differential expression of exons 2/3 at the N-terminal (Tapia-Rojas *et al*., 2019). In the early 90’s a high molecular weight tau isoform of tau was identified in adult peripheral nervous system (PNS) (Taleghany & Oblinger, 1992) and eventually cloned and termed Big tau (Couchie *et al*., 1992; Goedert *et al*., 1992; Georgieff *et al*., 1993). Big tau contains an additional exon (termed 4a) of 254/251 amino acids (in rat and human, respectively) expressed as a 110 kDa protein, increasing dramatically the length of the projecting domain relative to low molecular weight, (LMW) tau. In a comprehensive evolutionary analysis of MAPT gene and its exon structure with emphasis on Big tau (Fischer, 2022), the presence of the 4a exon was discovered at early stages of vertebrate evolution with a stable size of about 250aa but little or no sequence identity across species. This finding suggested that the 4a exon may have evolved independently in different species in response to the diversity of the evolving nervous system, generating a Big tau variant with a defined long projecting domain which increased the protein size and modified its structural and functional properties.

### 1.2. Big tau-early studies

The developmental expression of Big tau was studied in superior cervical ganglion (SCG) of the autonomic nervous system, and trigeminal ganglion and dorsal root ganglion(DRG) of the peripheral nervous system showing that it begins late in embryonic stages and gradually increases postnatally (Oblinger *et al*., 1991; Couchie *et al*., 1992; Goedert *et al*., 1992). In a recent study (Jin *et al*., 2023) we followed the regulation of Big tau during the development of SCG in more details and found that these neurons switch tau expression from the low molecular weight isoforms to Big tau, with the transition completed by about 4-5 weeks postnatally. We also confirmed the expression of Big tau in postnatal cultured SCG neurons during growth and in response to injury. Big tau is also expressed in spinal motor neurons, retinal ganglion neurons, the cerebellum and neurons that extend processes into the PNS, including most cranial nerve motor nuclei (Boyne *et al*., 1995). The analysis of Big tau so far has been focused on studies dealing mostly with distribution, structure and developmental regulation establishing the basic profile of Big tau. They did not however address most of the fundamental questions about the functional significance of the transition to Big tau. We speculated that Big tau facilitates axonal transport in projecting axons possible by increasing microtubule spacing (Fischer & Baas, 2020; Fischer, 2023). Similarly, the pathological significance of Big tau remained unknown, but interestingly neurons that express Big tau appear less vulnerable to tauopathies possible because this variant is less likely to form toxic aggregates.

### 1.3. Big tau in the CNS

There is also a gap of knowledge in the details of Big tau expression in selective CNS neurons such as the cerebellum and the visual system with the latter being the focus of the current study. The presence of Big tau in retinal ganglion cells (RGCs) was originally discovered in a study that explored the distribution of Big tau in the adult and developing nervous system (Taleghany & Oblinger, 1992; Boyne *et al*., 1995) showing that retinal ganglion cells and the optic nerve express Big tau at 90kDa, defined as middle molecular weight (MW) tau.

Subsequent studies (Mercken *et al*., 1995), which were focused on axonal transport of MAPs in RGCs of mice noted that the adult optic nerve contained both the LMW forms and the middle MW form of Big tau discovering that the transport rates of the tau proteins were distinct from that of tubulin and other MAPs. Interestingly, axonal transport rates in vivo (slow or fast) were unaffected by tau deletion or overexpression (Yuan *et al*., 2008) and did not show accumulations of vesicular organelles. The expression of tau in the visual system has also been studied in the context of microtubule dynamics, neuronal cell survival and regeneration as well as degeneration related to tauopathies and Alzheimer disease (AD). These studies have used a variety of experimental models including mouse models of AD and glaucoma, optic nerve crush, tau knockout and SiRNA-mediating silencing, and tau over expression. In addition, there has been significant progress in characterizing subpopulations of RGCs and discovering factors that promote survival and regeneration as well as the programs that control degeneration induced by injury (Duan *et al*., 2015; Aranda & Schmidt, 2021; Kim *et al*., 2021). Clinical studies contributed valuable knowledge from patients of AD and Parkinson’s Disease (PD) with findings about the effects of these diseases in the visual system relative to the molecular mechanism observed in brain. What has often not been addressed is the distinctive properties of tau in the visual system as one of the few CNS areas in which RGCs and the optic nerve also express the unique form of Big tau different from PNS neurons.

### 1.4. Big tau in the visual system

This study was designed elucidate the properties of Big tau in the visual system and in particular its unique isoform structure as a middle MW of 90kDa and its distribution in RGC and axons of the optic nerve. For the structural analysis we have focused our studies on testing 3 possible scenarios that may explain the 90kDa size: differences in alternative splicing of specific exons, changes in phosphorylation and the possibility of an initiation translational site downstream from the standard protein synthesis start codon at exon 1. For the distribution analysis we used Big tau specific antibodies in immunofluorescent staining experiments of RGCs and the optic nerve. We discovered that Big tau in RGCs expresses the 4a exon which defines Big tau, as well as exon 6 and exon 10 (one of the microtubule binding domains), but is lacking exons 2/3. Not surprising, Big tau share the extensive phosphorylation characteristic of tau isoforms present in other tissues. As far as the distribution of tau isoforms, it appears the RGCs and the optic nerve express both the LMW and the MMW isoforms throughout the neurons and their corresponding axons.

## 2. Results

### 2.1. MAPT

Fig 1A shows the general exon structure of MAPT including the N-terminal (N1 and N2 included in exons 2-3), the projection domain containing the proline rich area (exons 4-8) including exons 4a and 6 of Big tau, the microtubule binding domains (MTBD R1-R4 corresponding to exons 9-12) and the C-terminal (exons 13-14). In Fig. 1B we show the sequence of the first 4 exons. We will use these figures as a reference to our subsequent analyses.

**Fig. 1:**
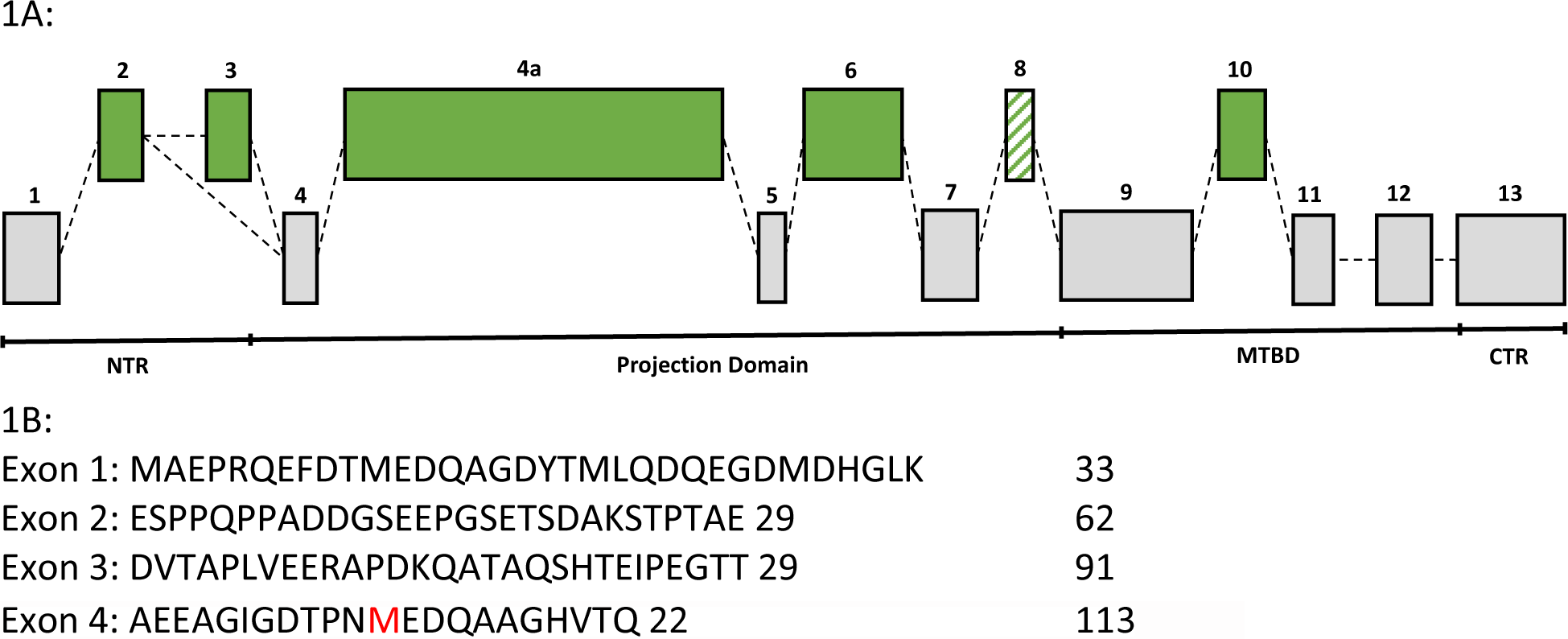
MAPT. Fig. 1A: Exon structure of MAPT showing 1) the different domains of tau including NTR, projecting domain, MTBD and CTR and 2) the different exons with alternative splicing including exons 2/3, 4a and 6 of Big tau and exon 10 that defines the 3R and 4R isoforms. Exon 8 which is hardly ever expressed is depicted in diagonal lines. NTR=N-terminal region. MTBD= microtubule binding domains. CTR= C-terminal region. 1B: Protein sequence of exons 1-4A of ENSRNOT00000042984.6 Mapt-207 (rat tau)

### 2.2. Analysis of exon structure

Fig. 2 show the analysis of Big tau exon structure in RGCs. Panel A.1 shows PCR analysis with primers from exon 2 and 4a resulting in a 923bp band in lanes 1 and 2 corresponding to the presence of both exons 2 and 3 in Big tau of DRG samples. This band is missing in lanes 3 and 4 of retina samples indicating the absence of exon 2. Since exon 3 is always expressed in combination of exon 2 while exon 2 can be expressed by itself (Himmler *et al*., 1989) it indicates the absence of both exon 2 and 3 in Big tau of the visual system. Panel A.2 shows PCR analysis with primers from exons 4A and 6 resulting in a band of 528bp in all samples (DRG lane 1 and 2 and retina lane 3 and 4) indicating the presence of exon 6 in all these variants of Big tau. Panel A.3 shows PCR analysis with primers from exons 9 and 12 with a single band of 306bp for retina samples in lane 3 and 4 corresponding to a 4R isoform only. In contrast, the DRG samples in lanes 1 and 2 show 2 bands at 212bp and 306bp corresponding to both 3R and 4R isoforms (3R= alternative splicing of exon 10 at 94bp).

**Fig. 2:**
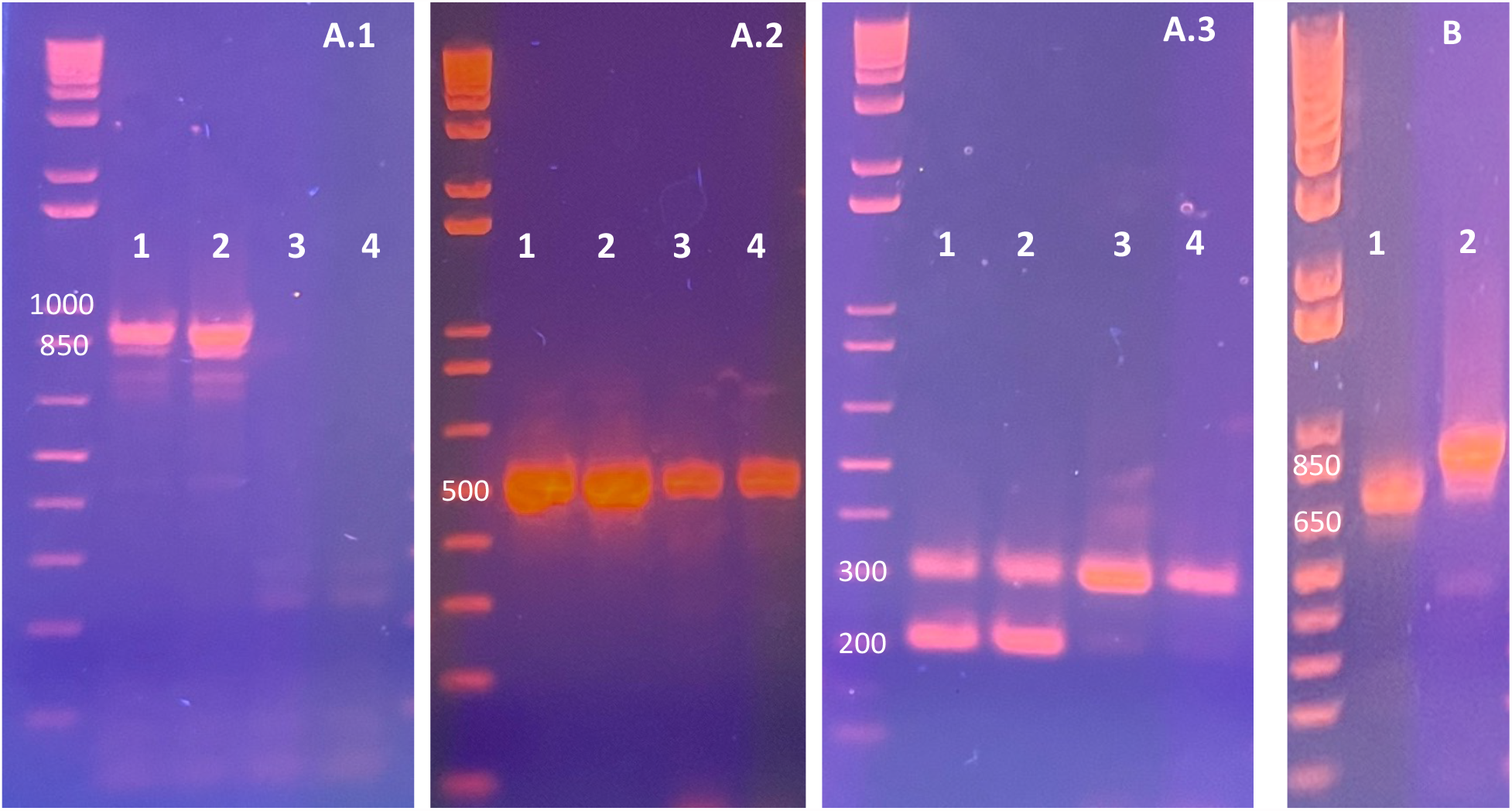
PCR analysis of Big tau exon structure in the visual system. A.1 exon 2 (5’ – TCA GAA CCA GGG TCG GA – 3’) exon 4A (5’ – GCA GGT TGC TTG TCA GTT GG – 3’) 2+/3+ = 923bp. DRG (lane 1 & 2), Retina (lane 3 & 4) A.2 exon 4A (5’ – GGG TTC CAT CCC ACT TCC TG – 3’) exon 6 (5’ – GGT GGT TCA CCT GAT CCT GG – 3’) 4a+/6+ = 528bp. DRG (lane 1 & 2), Retina (lane 3 & 4) A.3 exon 9 (5’ – CCG TCT GCC AGT AAG CGC – 3’) exon 12 (5’ –CTA CCT GGC CAC CTC CTG GC – 3’) 10+ = 306bp, 10− = 212 bp. DRG (lane 1 & 2), Retina (lane 3 & 4). B. exon 1 (5’ – CCG CCA GGA GTT TGA CAC AA – 3’) exon 4A (5’ – TGG AAT GTG AAC TCA GGG GC – 3’) 2+/3+ = 891bp, 2−/3 − = 717bp. Retina (lane 1), DRG (lane 2).

Taken together we show that transcripts of Big tau expressed by RGC neurons 1) lack exons 2 and 3 at the N-terminal (in contrast to Big tag tau in DRG and SCG), 2) Exons 4A and 6 are present all Big tau variants, 3) The dominant isoform of tau in the retina has the 4R domain with exon 10 (both in LMW and Big Tau). Although the lack of exons 2/3 in RGC neurons is likely to reduce the apparent MW of the Big tau protein relative to DRG and SCG neurons, it is not clear whether a reduction of 58 aa (Fig. 1B) is sufficient to explain the shift from 110kDa to 90kDa on Western blots, a 20% decrease that can be calculated to require about 100 aa. We posit that it possible that Big tau of the visual system is using a different translational initiation site from the original exon 1. Indeed, a second initiation site is present in exon 4 (Fig. 1B at 103aa from the primary initiation site, red) with a strong Kozak sequence (A at -3 and G at +4), which will generate a Big tau isoform 102aa shorter than the “conventional” Big tau of 110kDa (Fig xx).

We therefore designed primers starting at exons 1 and proceeded with a PCR analysis of exon 1 to 4A. The results show (Fig. 2B) that the visual system (lane 1) generated the predicted band that is smaller than the typical Big tau from DRG (lane 2) in the absence of exons 2/3. The analysis indicated that exon 1 is present in Big tau of the visual system and therefore excluded a second initiation site on exon 4.

### 2.3 Analysis of Big tau structure using Western blots

Fig. 3A shows a Western blot probed with Big tau antibodies comparing the retina/optic nerve with DRG and SCG. As previous studies have shown, the results confirmed that while Big tau of DRG and SCG displayed a protein at 110kDa, in the visual system Big tau has a MW of 90kDa often referred to as MMW tau. Fig. 3B/C shows the results using the tau-5’ and tau-3’ antibodies, respectively, confirming the MW of Big tau at110kDa and 90kDa for samples of DRG/SCG and visual system, respectively. They also reveal that the primary tau isoforms in the visual system are the LMW tau at 40-50kDa. The absence of exons 2/3 in the visual system means that tau-5’ antibodies recognized only 55aa instead of 112aa but are still likely to detect the tau isoforms. If, however, Big tau in the visual system is initiated from a second location on exon 4 there would have been only 11aa of reactivity, likely insufficient for detection. Fig. 3D shows the shift in the apparent MW of Big tau following dephosphorylation in DRG, SCG and samples from the visual system suggesting extensive phosphorylation in all samples.

**Fig. 3:**
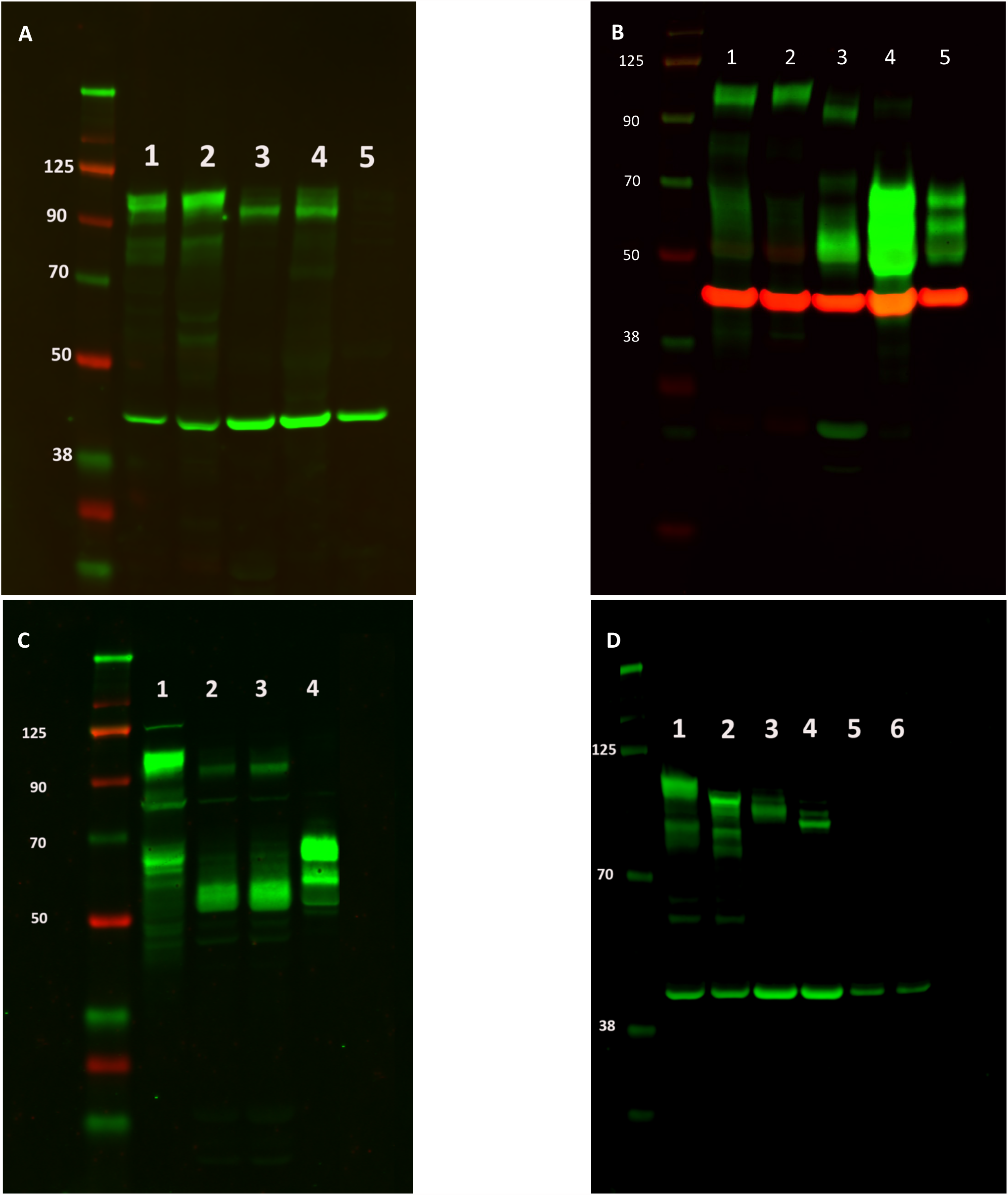
Western blot analysis of Big tau. Panel A: Big Tau antibody, lanes 1=SCG, 2=DRG, 3=Retina, 4=Retina 2X conc., 5=Brain Panel B: 3’ Tau antibody, lanes 1= DRG 2x conc., 2=DRG, 3=Retina, 4= Brain Panel C: 5’ Tau antibody, lanes 1= DRG, 2=Retina, 3= Retina 2X conc., 4= Brain 5X conc., 5=Brain Panel D: Big Tau antibody, lanes 1=DRG, 2-DRG dephos, 3=Retina, 4= Retina dehpos, 5= Brain, 6= Brain dephos.

Analysis of the developmental regulation of tau expression in the visual system included samples from E18, P5 and adult in comparison to embryonic and adult SCG (fig. 4). The results of Western blots indicated that at the embryonic stage (lane 2) only LMW tau could be detected at low levels (likely 3R), which increased dramatically at P5 (lane 3) with expression of higher molecular weight isoforms as well as a band at 80kDa which could be the MMW form of Big tau (90kDa) without exon 10 (3R). The adult retina (lane 4) showed the expected combination of LMW isoform together with the MMW Big tau (both 4R) while adult SCG (lane 6) showed the 110kDa isoform of Big tau consistent with our previous results (Fischer, 2023). The putative transition from 3R to 4R typical of tau maturation in general seems to manifest here as the gradual increase in MW in both LMW and Big tau isoforms shown when comparing lane 2 (E18) to lane 3 (P5) and then lane 4 (adult).

**Fig. 4:**
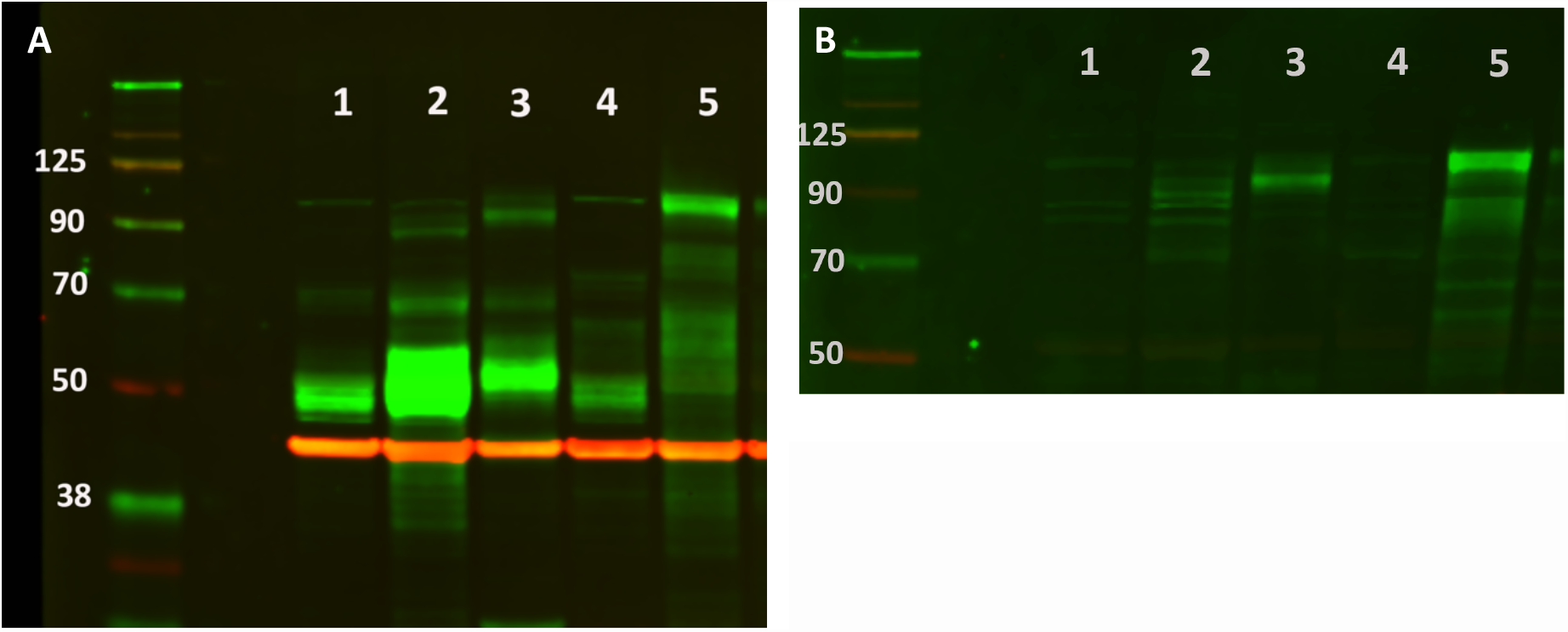
Developmental regulation of tau expression. Panel A: 3’ antibody. Panel B: Big tau antibody. Lanes: 1-whole eye E18, 2-Retina E18, 3-Retina P5, 4-Retina adult, 5-SCG.

### 2.4. Distribution of Big tau in the visual system

Fig. 5A shows double staining of the retina (pane A-C) and the optic nerve (panel D-F) with Big tau antibodies and Tuj. The results suggest that all RGCs and axon express the Big tau isoform. Comparable analysis with whole mount retina preparation shows similar results with Big tau antibodies staining all axons (Fig. 5B). It is however important to consider that the LMW isoforms of tau are the dominant variant in RGCs the optic nerve (see Western blots Fig. 3) and are likely to have the same wide distribution as Big tau.

**Fig 5:**
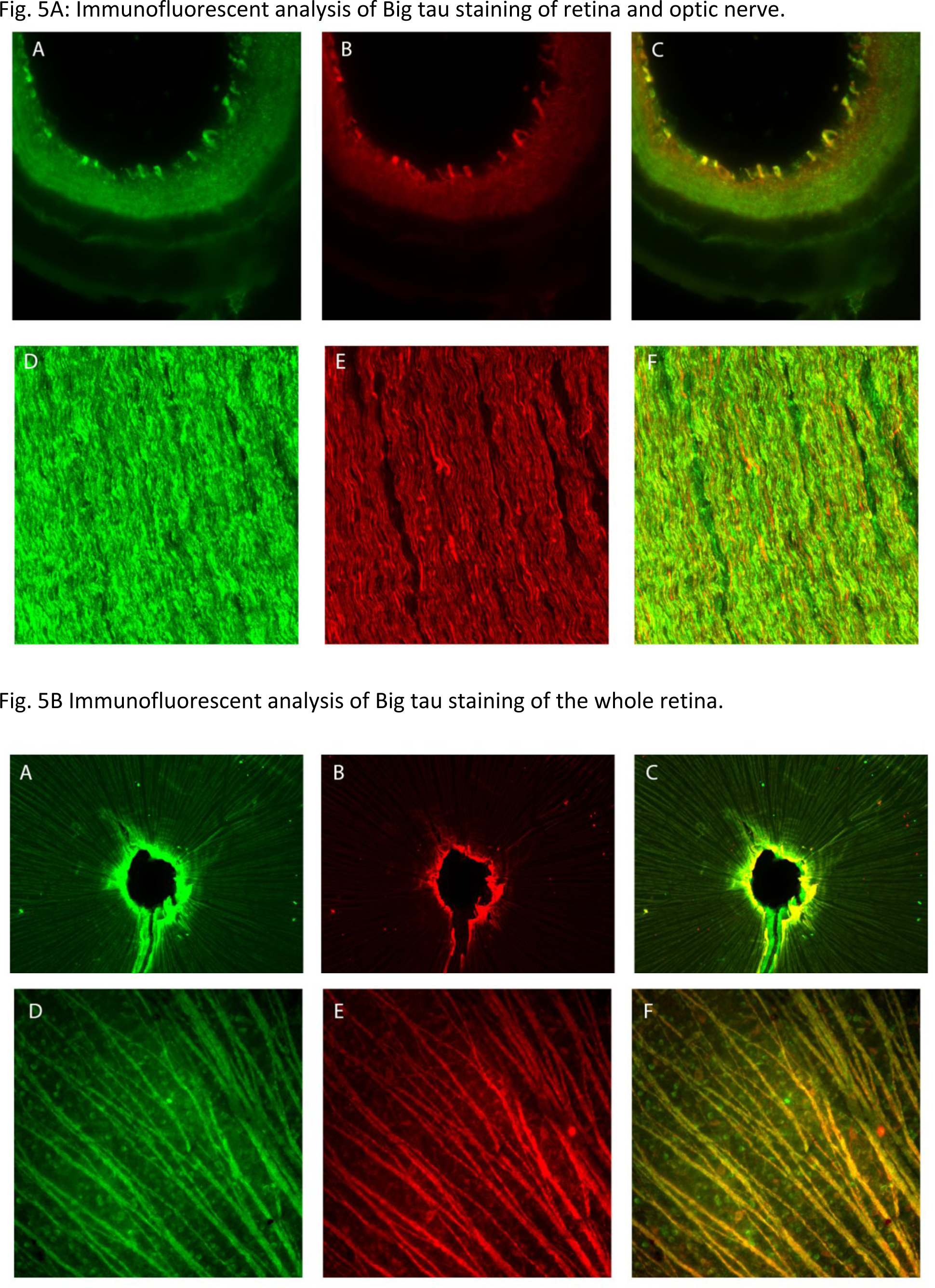
5A: Double staining of Big tau and Tuj for retina and optic nerve. Images show that retina coronal sections (A-C) and optic nerve sagittal sections (D-F) express Big tau (green, A and D) also express Tuj (red, B and E as well as C and F merged). Scale bar=1000 μm. 5B: Double staining of Big tau and Tuj for the whole retina. Images show the whole retina stained with Big tau (A) and Tuj (B) in low and Big tau (D) and Tuj (E) in high magnification. Most Tuj expressions were co-localized with Big tau (C and F) in low and high magnification. Scale =1000 μm.

## 3. Discussion

### 3.1. Major findings

After the original discovery and cloning of Big tau in the early 90’s (Couchie *et al*., 1992; Goedert *et al*., 1992), there has been a long pause in the study of Big tau properties. It was only in the last few years that the interest resumed with reports on its developmental regulation and evolutionary history (Fischer, 2022; Jin *et al*., 2023) as well as discovering the expression of Big tau in non-neural peripheral tissue such as Heart (Luciani *et al*., 2023), lung (Choi *et al*., 2022) and submandibular gland (Hamsafar *et al*., 2023) with implications on tau aggregation and function relative to LMW tau (Fischer & Baas, 2020; Fischer, 2023). Here we focused our attention to the visual system, considering early studies that reported a form of Big tau with a middle MW (Mercken *et al*., 1995), with the goal of elucidating the structure and expression in a complex but well define CNS system.

We report that the Big tau variant of the visual system is a protein of 90kDa which expresses 4a exon (defining Big tau) as well as exon 6 (like other forms of Big tau), but lacking exons 2/3. It also expresses exon 10 which determines the numbers of microtubule binding repeats (4R, typical of adult tau). The lack of exons 2/3 at the N-terminal is a common feature of the alternative splicing variations defining the 6 isoforms of LMW tau, but so far unique for Big tau. The absence of these exons may affect interactions with other proteins and likely responsible for the reduced MW of 90kDa. A preliminary analysis suggests that Big tau in the visual system is sharing the extensive phosphorylation characteristic of other tau isoforms. What is unique however, is that the adult visual system expresses both the low isoforms of tau and middle MW isoforms of Big tau (expressed in the retinal ganglion neurons and their corresponding axons). When the developmental regulation of tau expression was examined, we found that at embryonic stages low levels of only LMW tau are expressed increasing dramatically at early postnatal stages with adult retina expressing the LMW of tau together with the MMW isoform of Big tau. It appears that during development there is a transition from 3R to 4R isoforms in both LMW tau and Big tau. Finally, we report that Big tau and likely the LMW isoforms of tau appear to be expressed in all RGC and their exons.

### 3.2. Limitations of the study

a) Although we have strong evidence for the combined expression of Big tau and LMW in adult visual system using Western blots and the presence of Big tau in all RGCs and their axons using Big-tau specific antibodies, we can only assume that a similar distribution occurs for LMW tau since there are no antibodies for LMW tau only. The final analysis could be accomplished by reviewing data of single cell sequences. b) The absence of exons 2/3 translates into a reduction of 58aa which may not be sufficient to result in a shift in the of 110kDa to 90kDa (theoretically it needs >100aa). We did not find size changes in exons 1 and 4, but it is possible that other exons are shorter and contribute to the reduction in the apparent MW. We did present a general assessment of a highly phosphorylated Big tau protein but do not have data to examine changes in other post-translational modification that could affect Big tau structure. c) We speculate about the functional implications of the unique structure and expression of Big tau but do not have experiment to test them.

### 3.3. Conclusions

The story of Big tau is getting increasingly complex and interesting. In the early days of Big tau discovery (Georgieff *et al*., 1991; Goedert *et al*., 1992; Boyne *et al*., 1995), it appeared to be expressed mostly in peripheral neurons setting a distinction between the PNS and CNS systems. It led to a proposal on a putative function in long peripheral axons that need efficient axonal transport and support against metabolic stress. The dramatic increase in the size of the projecting domain of tau with the addition of exon 4a was consistent with a potential increase in microtubule spacing or related mechanisms that can facilitate axonal transport (Fischer & Baas, 2020). The structural changes in Big tau could also affect the pathological aggregation of the protein providing protection against tauopathies. Consistent with these ideas was evidence from an evolutionary study (Fischer, 2022) that showed that despite sequence diversity of the 4a exon along the vertebrate lineages, its size of about 250 aa was stable. There is also clinical evidence of relatively more resistance or delay of neurodegeneration of PNS neurons relative to CNS neurons. Things started to get more complex when even in PNS the expression of Big tau in DRG neurons was shown to be selective to small and medium size cells and the finding that at early developmental stages SCG neurons expressed only the LMW forms and then switch to express Big tau postnatally (Jin *et al*., 2023). Further analysis discovered that in the CNS there are specific regions such as the cerebellum and the visual system that express Big tau (Boyne *et al*., 1995). Here we report another level of complexity in Big tau expression with a variant that lacks exons 2/3 leading to a middle MW isoform of Big tau of 90kDa and the co-expression of both the LMW tau and Big tau in adult RGC and optic nerve. In addition, it is possible that at early postnatal stages it is expressed as a 3R isoform at 80kDa before the transition to the adult 4R isoform at 90kDa. These data suggest that Big tau has a variety of functions either alone or in combination with other isoforms of LMW tau. Thus, it is possible that the co-expression of Big tau with the LMW isoforms of tau (with relatively low levels of Big tau/LMW tau) impedes its protection capacity against aggregation leading to neurodegeneration. Indeed, clinical data indicate that AD is associated with visual impairment, optic neuropathy and RGC loss which occurs by Aβ deposits and phosphorylated tau generating aggregates (Gasparini *et al*., 2011; Ho *et al*., 2012; Hart *et al*., 2016; Romaus-Sanjurjo *et al*., 2022). Given the heterogeneity of RGC with respect to distinct subtypes (Duan *et al*., 2015; Aranda & Schmidt, 2021; Kim *et al*., 2021) a selective expression of Big tau could have provided an interesting correlation with properties of individual neurons or type of neurons. Our finding of Big tau expression in all RGCs and their axons excludes a direct correlation with improved regeneration capacity or survival following injury. Indeed, previous studies indicated that the deletion of tau did not affect regeneration and RGC survival in the injured mouse visual system (Rodriguez *et al*., 2020) or the rate of axonal transport (Yuan *et al*., 2008). Such a conclusion is also consistent with regeneration experiments of the sciatic nerve showing that the levels of Big tau protein decreased in the DRG following axotomy, without a change in the pattern of the other tau isoform, a reduction that was mirrored with a reduction in the levels of Big tau mRNA (Oblinger *et al*., 1991). The enigmatic properties of Big tau in the visual system add another dimension to the remaining mysteries of its function (Fischer & Baas, 2020) and distinct roles in the PNS and CNS system. As a reminder, the cerebellum is the other unexplored CNS region expressing Big tau that may reveal more surprises. At any case, with increased data on Big tau structure and expression in a variety of systems, the stage is ready for more direct functional experiments in either in vitro systems that express distinct isoforms of tau or in vivo by gene deletion and/or over expression. Understanding the functional properties of Big tau may also open a therapeutic window in the neurodegeneration field particularly related to tauopathies.

## 4. Methods

### 4.1. Antibodies

Fig. 6 shows the tau antibodies we prepared against recombinant proteins of tau as previously described (Boyne *et al*., 1995; Black *et al*., 1996). They include the N-terminal (112aa, tau 5’ Ab), the C-terminal (72aa, tau 3’ Ab) and the complete 4A exon of Big tau (254aa, Big tau Ab). These antibodies will be used for both Western blot analysis and immunofluorescence staining.

**Fig. 6:**
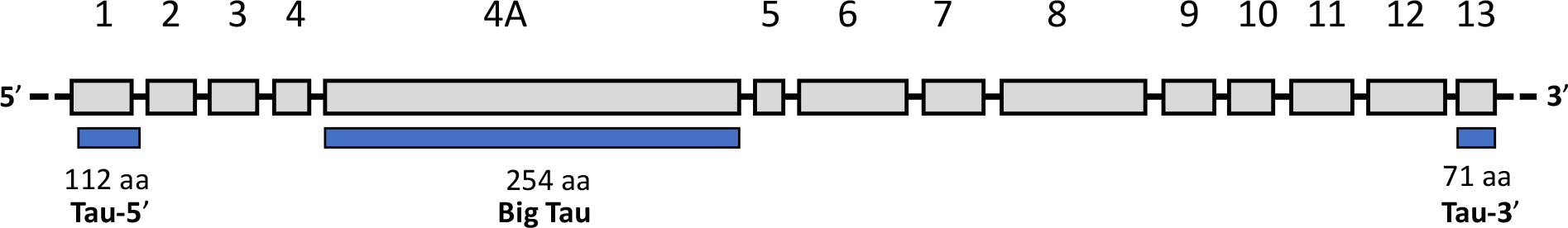
Tau antibodies. Tau 5’ antibodies were prepared against 112aa at the N-terminal (2-113, exon 1-4) Tau 3’ antibodies were prepared against 71aa at the C-terminal (618-686aa exon 13-14) Big tau antibodies were prepared against the complete 254aa of the 4A exon.

### 4.2. Tissue preparation

Retina and optic nerve were dissected from adult rats after perfused with 0.9% saline and 4% paraformaldehyde (PFA) and kept in 4% PFA overnight. Retina and optic nerve were transferred to 30% sucrose for 3-4 days. For tissue section, retina was cut coronally, and optic nerve was cut sagittally at 20um and mounted on the slides, and stored at -20° C for further staining. For whole retina, after incubated in 4% PFA overnight, tissue was transferred to PBS, stored at 4° C.

### 4.3. Immunohistochemical staining

For section staining, slides with retina and optic nerve were washed with phosphate-buffered saline (PBS) for 3 times and incubated in blocking solution (PBS with 0.3% triton X-100, 0.5mg/ml BSA, 0.01% thimerosal, 5-10% goat serum) for 1 hour at room temperature. Sections were incubated in the following primary antibodies overnight at room temperature for double staining: Big tau (1:5000, Fischer lab) with Tuj (1:1000, Covance). Sections were incubated in the secondary antibodies: goat anti-rabbit AF 488 (1:1000, Jackson ImmunoResearch), goat anti-mouse AF 594 (1:1000, Jackson ImmunoResearch) for 2 hours at room temperature after washing in PBS for 3 times. Sections were coverslipped with anti-faded solution.

For the whole retina staining, the whole retina was washed with PBS for 3 times in 48-well plate. Retina was blocked with blocking solution same as above for 1 hour at room temperature. Retina was incubated in the primary antibodies same as section staining overnight at room temperature. On next day, sections were incubated in the secondary antibodies: goat anti-rabbit AF 488, goat anti-mouse AF 594 for 2 hours at room temperature after washing in PBS for 3 times. Retina was mounted on the slide and coverslipped with anti-faded solution.

### 4.4. Western blot analysis

#### 4.4.1. Tissue preparation

Tissue homogenates derived from rat neural tissues were disrupted and homogenized in cold lysate buffer (RIPA lysis buffer: 25 mM Tris-HCl pH 7.6, 150 mM NaCl, 1% NP-40 or Triton X-100, 1% sodium deoxycholate, 0.1% SDS) in the presence of protease and phosphatase inhibitors using a Fisher brand Sonic Dismembrator. Protein determination was done by Pierce™ BCA Protein Assay Kit (Thermo Fisher). The tissue homogenates were stored at −80°C. For dephosphorylation tissue samples were homogenized in extraction buffer without phosphatase inhibitors or detergent as described above and protein determinations were made. Duplicate aliquots of each sample were prepared (one for dephosphorylation and one for control). New England BioLabs Lambda Protein Phosphatase Kit was used to prepare dephosphorylated samples. All samples were stored at −80°C and prepared for Western blotting described below.

#### 4.4.2. SDS gels

Equal amounts of protein from each sample were denatured in loading buffer containing DTT by boiling at 95°C for 5 min. Protein samples and Chameleon duo pre-stained protein ladder (LI-COR) were separated by sodium dodecyl sulfate polyacrylamide gel electrophoresis (SDS-PAGE) on 4%–12% polyacrylamide bis-tris gels (Thermo Fischer). After distilled water washing and dry transfer (Thermo Fischer iBlot system), PVDF membranes were washed in TBS and blocked in Intercept Blocking Buffer (LICOR) for 1 h at room temperature, followed by overnight incubation in primary antibody at 4°C. Rabbit polyclonal primary antibodies to 3’and high MW Tau were prepared in-house and used at 1:5K and 1:1K respectively. All primary and secondary antibodies were diluted in Intercept Antibody Diluent (LICOR). Membranes were washed with TBS-Tween 20 and incubated with fluorescently labeled IRDye secondary antibodies (LI-COR) for 1h at room temperature. After TBS washes, membranes were washed in distilled water and digital fluorescent visualization of signals was detected at 700 and/or 800 nm channels using the Odyssey^®^ CLx Imaging System (LICOR).

### 4.5. Polymerase chain reaction (PCR)

The extraction and purification of adult Sprague-Dawley rat RNA was performed using a RNeasy Mini Kit (Qiagen, Germantown, MD). cDNA was obtained using High-Capacity cDNA Reverse Transcription kit (Applied Biosystems, ref. 4368814, Thermo Fisher Scientifics, Waltham, MA).

PrimerBLAST (BLAST [internet National Library of Medicine],Bethesda, MD:) was used to design primers to target specific exons as followed: exon 1 (5’ – CCG CCA GGA GTT TGA CAC AA – 3’) and exon 4A (5’ – TGG AAT GTG AAC TCA GGG GC – 3’), exon 2 (5’ – TCA GAA CCA GGG TCG GA – 3’) and exon 4A (5’ – GCA GGT TGC TTG TCA GTT GG – 3’), exon 4 (5’ – GCA GGC ATC GGA GAC ACC CCG A – 3’) to exon 7 (5’ – TGT GGC GAT CTT CGC CCC CGT T – 3’); exon 4A (5’ – GGG TTC CAT CCC ACT TCC TG – 3’) to exon 6 (5’ – GGT GGT TCA CCT GAT CCT GG – 3’); and exon 9 (5’ – CCG TCT GCC AGT AAG CGC – 3’) to exon 12 (5’ –CTA CCT GGC CAC CTC CTG GC – 3’ All primers were purchased from IDT (Integrated DNA Technologies, Inc, Coralville, IA). PCR were run using a Taq DNA polymerases Kit (ABM, Richmond, B.C) and a thermal cycler (Bio-Rad, Hercules, CA). Electrophoresis was performed on 1.5% agarose gel at 94V for 2 hours.

## Acknowledgment

We acknowledge the core facility of the Marion Murray Spinal Cord Research Center at the Department of Neurobiology and Anatomy. This work was supported by the William P. Snyder, III Chair Endowment to IF.

## Author contributions

YJ – performed the immunofluorescent staining, TC performed the Western blot analysis, JB performed the PCR experiments and prepared the figures, IF designed the experiments and wrote the paper. All authors agreed on the final draft of the paper.

## Sequence of MAPT

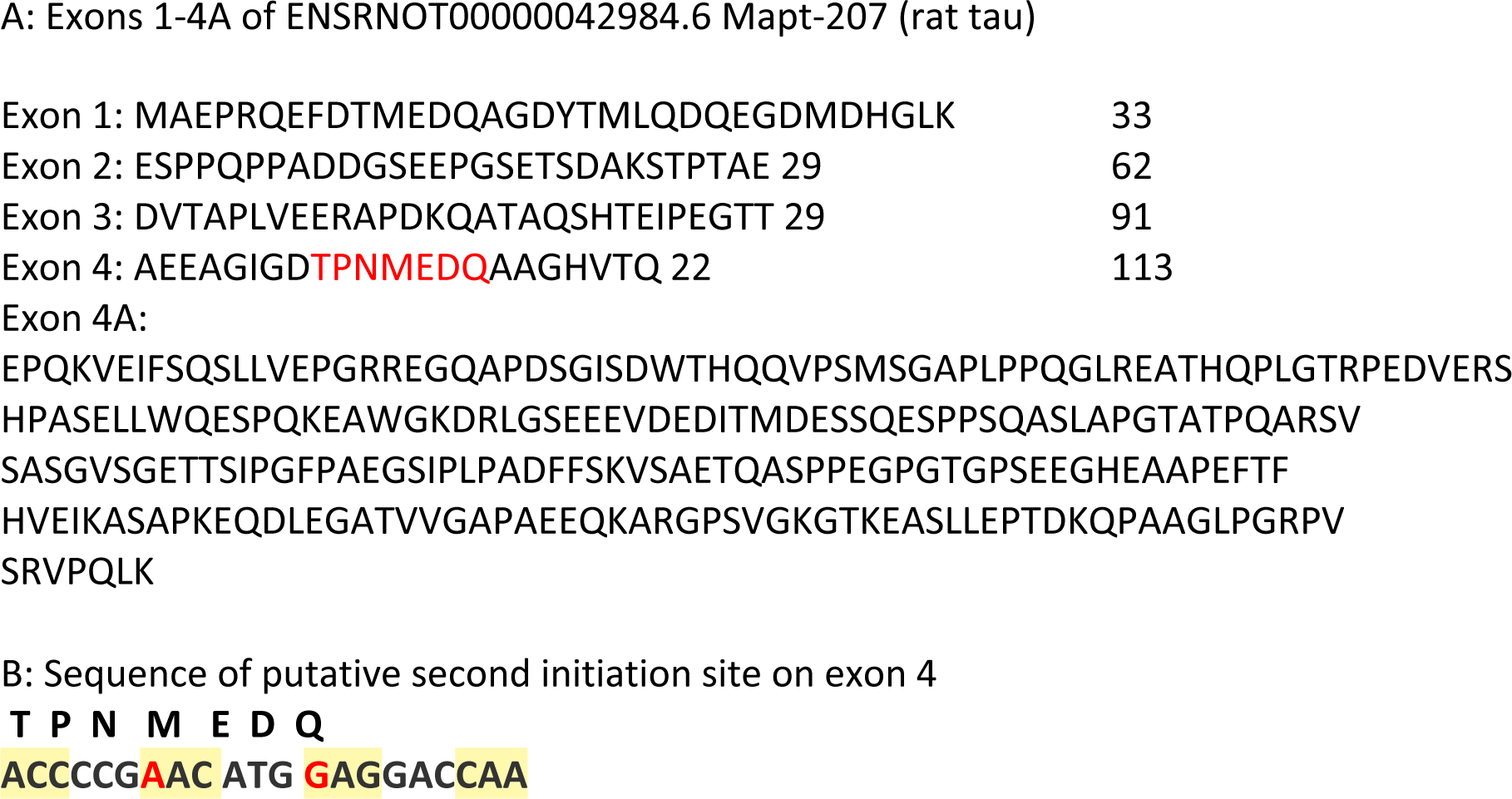

